# Antibiotics and copper drive compartment-specific dysbiosis and functional reprogramming in tomato microbiomes

**DOI:** 10.64898/2026.05.27.728321

**Authors:** Fabiano José Perina, Vanessa Thomas, Toi Ketehouli, Sameerika D. Mudiyanselage, Erica M. Goss, Samuel J. Martins

## Abstract

Plant-associated microbiomes help sustain plant immunity and productivity, yet the degree to which agricultural antimicrobials reshape these microbial networks and alter disease outcomes remains poorly characterized. Here, we quantified how chemical inputs such as copper and streptomycin trigger distinct, compartment-specific dysbiosis in tomato (*Solanum lycopersicum*), fundamentally decoupling microbiome-mediated immunity from pathogen defense. We demonstrated that chemical disturbance correlates with increased susceptibility to bacterial spot caused by *Xanthomonas perforans*. Notably, streptomycin-induced dysbiosis increased epidemic intensity, characterized by physiological and growth trade-offs, including reduced fruit number, mass, and seed weight, alongside a decline in photosynthetic gas exchange; while copper-induced dysbiosis had intermediate effects. Chemical perturbation restructured microbiomes in a niche-dependent manner based on amplicon profiling: seeds and the phyllosphere showed the greatest instability, including higher dispersion, taxon turnover, and network reorganization under streptomycin, whereas rhizosphere communities remained more deterministic but were structurally reshaped by copper. Rhizosphere metagenomics further revealed enrichment of antibiotic-resistance functions under streptomycin treatment and of metal-tolerance and oxidative-stress functions under copper treatment, along with shifts in genes linked to cell-envelope remodeling and redox metabolism. These findings identify microbiome dysbiosis as a mechanistic bridge between chemical stress and plant disease, and support crop protection strategies that preserve microbiome integrity.

## Introduction

Plants are associated with diverse and dynamic microbiomes that occupy distinct ecological niches throughout the plant, including belowground and aboveground compartments such as the rhizosphere, endosphere, phyllosphere, flowers and seeds. The microbial community is shaped not only through the environment, such as soil, water, and air^1, 2^. Collectively, microbial communities are essential to plant performance and are regarded as a “second genome” due to their critical roles in plant immunity system in addition to nutrient absorption, phytohormone regulation, and stress tolerance^3, 4^. The loss or disruption of beneficial microbiota can impair important host-associated functions, resulting in a state of dysbiosis^5, 6^. Dysbiosis may arise from abiotic stressors, such as antibiotic exposure or heavy metal contamination, as well as biotic pressures, including pathogen infection and disease progression^7, 8, 9^.

The control of bacterial plant diseases in modern agriculture relies heavily on chemical inputs, including copper-based bactericides and, in some cases, antibiotics such as streptomycin and oxytetracycline^10^. Although these compounds can effectively suppress phytopathogens, their persistent use has raised concerns regarding unintended ecological consequences within soils and plant-associated microbiomes. Chemical stressors, particularly antibiotics and heavy metals such as copper-based compounds, can persist and accumulate in agricultural soils, exerting strong selective pressures that reshape microbial community structure and function^6, 11, 12, 13^. These disturbances can promote the evolution of resistance determinants, horizontal gene transfer, detoxification pathways, oxidative stress responses, and efflux mechanisms^14, 15, 16, 17^. Beyond altering taxonomic composition, such stressors may reprogram microbial functional capacity, thereby modifying microbiome contributions to nutrient cycling, immune priming, niche occupation, and overall plant health. Importantly, these effects may decouple microbiome structure from function, resulting in communities that appear taxonomically altered yet functionally resilient, or conversely, functionally compromised despite relatively minor compositional changes^15, 16^. Despite these recognized effects, their integrated consequences across plant-associated microbiomes and subsequent effects on plant health remain inadequately understood.

Growing evidence suggests that microbial communities exhibit significant variation across plant compartments and that disturbances may have compartment-specific impacts^18, 19^. Furthermore, plant microbiomes are not confined to a single generation; seed-associated microbial communities may serve as reservoirs for vertical transmission, potentially perpetuating dysbiotic conditions across plant generations^20, 21^. In this study we hypothesized that antibiotic and copper applications cause systemic microbiome dysbiosis across plant compartments, including the rhizosphere, endosphere, and seeds, thereby directly affecting plant health and disease vulnerability. We propose that (i) antibiotic and copper treated plants demonstrate modified microbial community composition and functional capacity in these compartments, resulting in heightened vulnerability to infection by *Xanthomonas perforans*, and (ii) antibiotic exposure influences host responses, culminating in increased expression of plant defense-related genes during pathogen challenge.

To test these hypotheses, we conducted two greenhouse experiments in tomato (*Solanum lycopersicum*) using three treatments: sterile distilled water, copper hydroxide, or streptomycin to induce microbiome dysbiosis. Treatments were first applied to the rhizosphere as a soil drench at 21 days after emergence, and plants were challenged with *Xanthomonas perforans* (syn. *X. euvesicatoria* pv. *perforans*) 24 h later. The *X. perforans* strain used in this study was selected because it exhibited tolerance to both copper and streptomycin. A second application at flowering targeted both the rhizosphere and phyllosphere via an additional soil drench and foliar spray. We combined disease, physiological, and yield assessments with compartment-resolved 16S rRNA gene sequence profiling of the rhizosphere, phyllosphere, and seeds, along with rhizosphere shotgun metagenomics, to define how agricultural antimicrobials reconfigure microbiome structure and function across plant compartments. By linking chemically induced dysbiosis to pathogen susceptibility and host performance, this study establishes a mechanistic framework showing how antimicrobial inputs destabilize plant-associated microbiomes and identifies microbiome integrity as a critical, yet overlooked, dimension of sustainable disease management.

The objectives of this study were to: (i) determine how copper hydroxide and streptomycin applications alter microbial community composition across the rhizosphere, phyllosphere, and seed compartments of tomato plants; (ii) evaluate the effects of chemically induced microbiome dysbiosis on plant health and susceptibility to infection by *X. perforans*; (iii) characterize functional changes associated with dysbiosis using rhizosphere shotgun metagenomics and compartment-resolved 16S rRNA gene profiling; (iv) assess plant physiological, growth, and defense-related responses following antimicrobial exposure and pathogen challenge.

## Results

### Impact of dysbiosis on disease severity, yield, and physiology

To evaluate whether antimicrobial-induced shifts in microbiome structure alter disease outcome, we quantified the progression of bacterial spot caused by *X. perforans* following post-infection and application of copper and streptomycin. Cumulative disease severity differed markedly among treatments (Fig. 1A), as quantified by AUDPC (area under the disease progress curve). Antimicrobial treatments significantly increased bacterial spot development, with streptomycin-treated plants exhibiting a 54% higher AUDPC than the control (p < 0.001), followed by copper-treated plants, which showed a 21% increase (p < 0.01). These results indicate a high treatment-dependent amplification of epidemic intensity, with streptomycin applications producing the greatest cumulative disease burden.

**Fig. 1.**
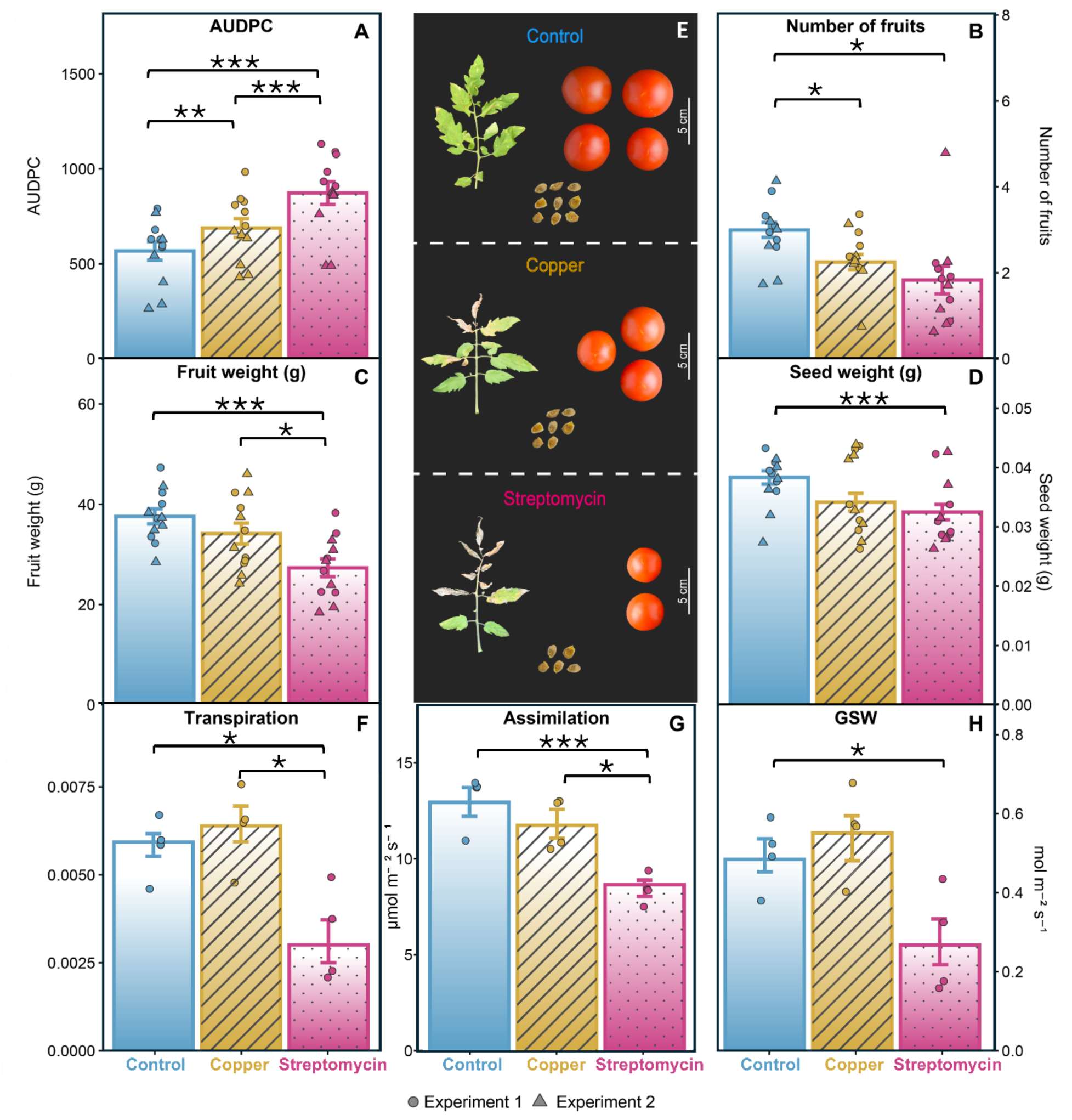
Effects of induced dysbiosis with copper [inline] and streptomycin [inline] on disease severity and yield traits in tomato plants. (A) Area under the disease progress curve (AUDPC) for bacterial spot caused by *Xanthomonas perforans*, assessed over seven evaluations from 5 to 23 days after inoculation (DAI) at 3-day intervals. Streptomycin-treated plants showed the highest disease severity, followed by copper-treated plants, relative to controls. (B) Number of fruits per plant harvested until 100 days after transplanting. Streptomycin- and copper-treated plants produced 39% and 25% fewer fruits than control plants, respectively. (C) Average fresh mass per fruit in grams. Streptomycin and copper treated plants showed reductions of 27% and 91%, respectively, relative to controls. (D) Average seed weight per plant, determined from 25 seeds per treatment and replicate. Streptomycin and copper-treated plants showed reductions of 12% and 6%, respectively, relative to controls. (E) Representative phenotypes for each treatment, showing (from top to bottom) leaves, fruits, and seeds from control (top), copper (middle), and streptomycin (bottom) plants. (F–H) Leaf gas-exchange parameters measured at 24 DAI: (f) Transpiration rate (mol m⁻² s⁻¹); (g) net CO₂ assimilation (µmol m⁻² s⁻¹) Streptomycin and copper-treated plants showed reductions of 21% and 10%, respectively, relative to controls; (h) Stomatal conductance to water vapour, gsw (mol m⁻² s⁻¹) Streptomycin-treated plants exhibited a 16% reduction in stomatal conductance relative to controls, while copper-treated plants showed an 13.4% increase. In all panels, significance annotations reflect Tukey’s HSD pairwise comparisons: *, p<0.05; **, p<0.01; ***, p<0.001; ns, non-significant. Bars represent treatment means ±SE (n = 6 plants per treatment per experiment). Overlaid points (circles, experiment 1; triangles, experiment 2) correspond to individual experimental units.

The temporal dynamics of disease severity closely paralleled the cumulative differences in disease severity. Disease progression diverged from the second assessment onward, with antimicrobial treated plants exhibiting faster symptom development than control plants between 5 and 23 days after inoculation at 3-day intervals (Supplementary Fig. S1A). From the third assessment onward, streptomycin-treated plants showed higher disease severity than controls, remaining significant through the final assessment. Streptomycin exceeded copper at assessments 5-7, while the control only surpassed copper at assessments 6-7. These temporal patterns mirrored the AUDPC ridgeline distributions (Supplementary Fig. S1B). Control plants exhibited lower disease severity distributions, whereas copper-treated plants showed an intermediate shift from low to high disease severity over time. In contrast, streptomycin-treated plants displayed the broadest distribution of disease severity and the greatest proportion of highly diseased plants throughout the experiment (Supplementary Fig. S1B). Corresponding to increased disease intensity, accelerated epidemic progression, and an expanded range of severity outcomes relative to control and copper-treated plants. Antimicrobial treatments also negatively affected yield. Streptomycin- and copper-treated plants produced fewer fruits, with reductions of 39% and 25%, respectively (Fig. 1B–E). Streptomycin-treated plants exhibited impaired resource allocation to progeny, with fruit and seed mass declining by 27% and 12%, respectively, relative to controls (Fig. 1F–H).

To determine the physiological basis of this host decline, we assessed photosynthetic performance under chemical stress using a LI-6800 portable photosynthesis system (Li-Cor, Lincoln, NE, USA). Physiological responses were affected by antimicrobial treatment only in Experiment 1. Streptomycin-treated plants exhibited significantly lower net CO₂ assimilation and transpiration compared to both control and copper-treated plants, whereas the latter two did not differ. Stomatal conductance (gsw) differed only between streptomycin and copper treatments, while intercellular CO₂ concentration (Ci) remained unchanged. In Experiment 2, no significant differences were detected for any physiological parameter.

### Chemical stressors differentially impact microbiome stability across plant compartments

To elucidate the ecological impacts of chemical exposure, we characterized bacterial community responses across the compartments: rhizosphere, phyllosphere and seeds using 16S rRNA gene amplicon sequencing. We found that microbiome restructuring was strongly governed by compartment-specific factors, reflecting differences in ecological filtering, community complexity, and resilience to disturbance. Although both chemical treatments altered community composition relative to controls, the magnitude and ecological signatures of these differences varied across plant compartments. Aboveground microbiomes (phyllosphere and seeds), which exhibited lower taxonomic richness, were more sensitive to chemical perturbation, with seed microbiome showing the greatest loss of core taxa, retaining only three OTUs representing 29% of total abundance across treatments. In contrast, rhizosphere communities displayed greater structural stability. These compartment-dependent responses were consistently supported across multiple analytical approaches.

### Seed microbiomes exhibit pronounced dysbiosis under streptomycin and copper stress

To determine the impact of chemical perturbation on the primary microbial reservoir, we characterized the seed-associated microbiota, the foundational inoculum for the developing plant. While the seed compartment exhibited the lowest OTU (Operational Taxonomic Unit) richness across all compartments, it maintained a conserved phylum architecture dominated by Proteobacteria, Firmicutes, and Actinobacteria across treatments. Despite this conserved phylum-level composition, community structure diverged among treatments. Control samples clustered tightly in NMDS space, indicating low compositional variability, whereas streptomycin-treated seeds showed the greatest dispersion, consistent with increased stochasticity and dysbiosis (Fig. 2C). Copper-treated communities also diverged from controls, but with lower dispersion than did streptomycin-treated communities. Bray-Curtis dissimilarities were highest under both stressors, while the beta nearest taxon index (*β*-NTI) which indicates the balance between deterministic and stochastic assembly processes, values near zero across treatments indicated predominantly stochastic assembly, likely reflecting low OTU richness. A minimal core microbiome was identified, with only three shared OTUs accounting for 29% of total abundance, and most taxa were treatment-specific. Linear discriminant analysis was used to specifically identify taxa most strongly associated with treatment groups, revealing enrichment of OTUs classified as *Xanthomonas* in streptomycin-treated seeds and *Pseudomonas* in controls. Network analysis (Fig. 2F) further revealed treatment-induced dysbiosis. Compared to control networks (82 nodes, 396 edges, 11 hubs), copper treatment caused a marked collapse (10 nodes, 46 edges, 3 hubs), while streptomycin resulted in a moderate reduction (59 nodes, 202 edges, 10 hubs). These declines in network size and connectivity indicate reduced community complexity and potential functional resilience under dysbiotic conditions.

**Fig. 2.**
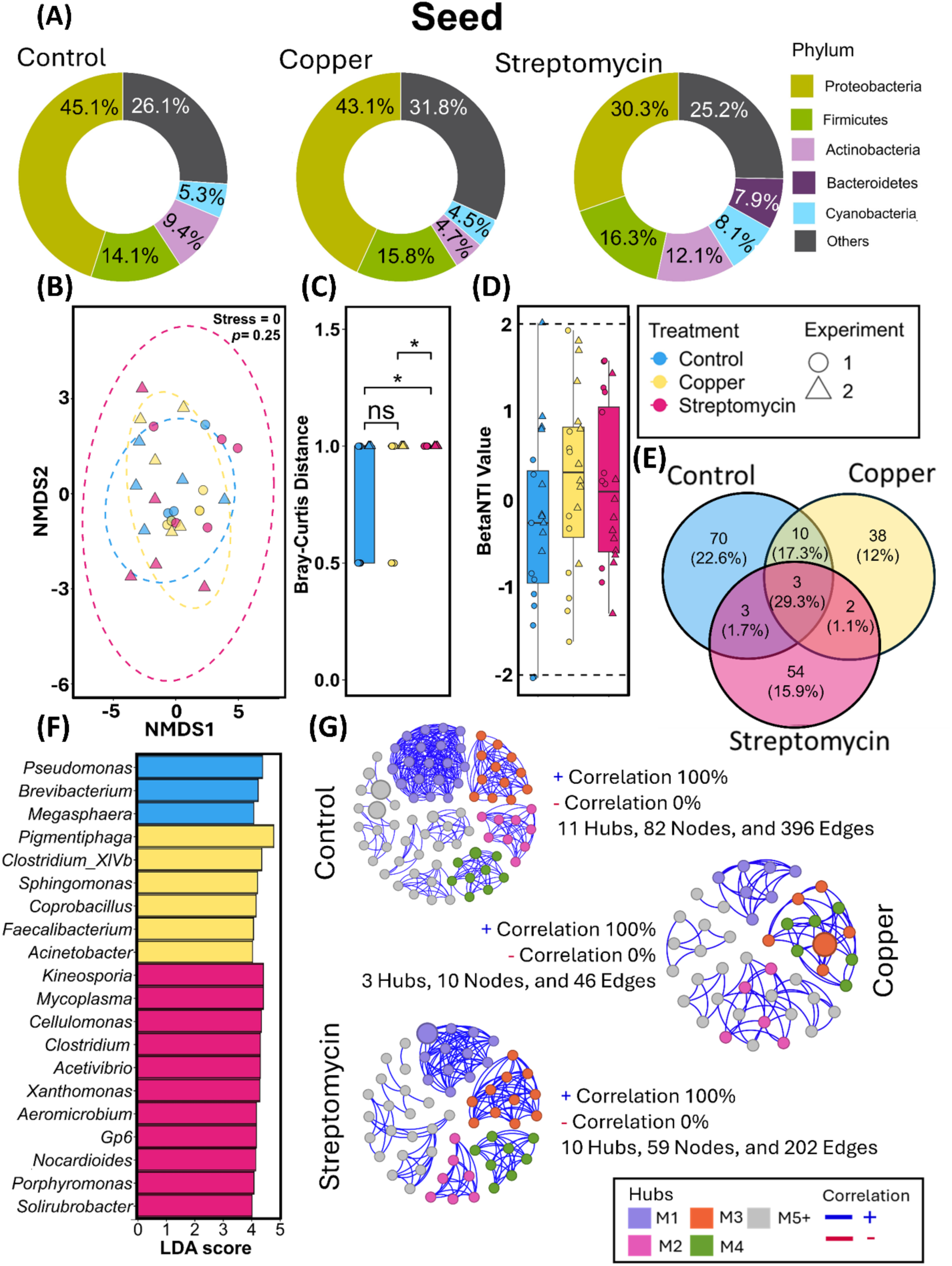
Seed microbial communities were analyzed for relative taxonomic composition at the phylum level using donut plots (A). Community composition similarity among streptomycin, copper, and control treatments was assessed using NMDS ordination based on Bray-Curtis dissimilarities (B), and Bray-curtis distance analysis was used to compare treatment separation (C). *β*-nearest taxon index (*β*-NTI) was used to evaluate microbial assembly processes and dysbiosis (D). Venn diagrams identified shared core microbiota, loss of core taxa, and treatment-specific taxa (E). Linear discriminant analysis (LDA) identified differentially enriched taxa among treatments (F), and Spearman co-occurrence networks were used to assess microbial interaction patterns (G).

### Streptomycin and copper destabilize phyllosphere community assembly despite conserved taxonomic composition

To investigate the impact of chemical perturbation on the leaf-surface microbiota, we characterized phyllosphere community dynamics using high-depth 16S rRNA gene sequencing. We demonstrate that the phyllosphere exhibited a significant predominance of Proteobacteria (>95% relative abundance) across all three treatments, indicating pronounced host and environmental filtering (Fig. 3A). Despite this taxonomic consistency, community structure varied substantially with treatment. Streptomycin-treated samples exhibited the greatest dispersion in NMDS space and the highest Bray-Curtis dissimilarities, reflecting increased stochasticity. Copper-treated communities formed the tightest clusters, consistent with stronger deterministic filtering. A large core microbiome comprising 813 shared OTUs accounted for 99% of total abundance; however, streptomycin treatment markedly increased taxon turnover, with over 200 unique OTUs detected exclusively under antibiotic exposure. *β*-NTI values further supported enhanced stochastic assembly in streptomycin-exposed communities, whereas control communities exhibited more structured assembly. Network analysis revealed predominantly positive associations across treatments, but streptomycin-treated communities displayed markedly expanded networks, suggesting proliferation of treatment-tolerant co-occurring taxa rather than recovery of stable community structure.

**Fig. 3.**
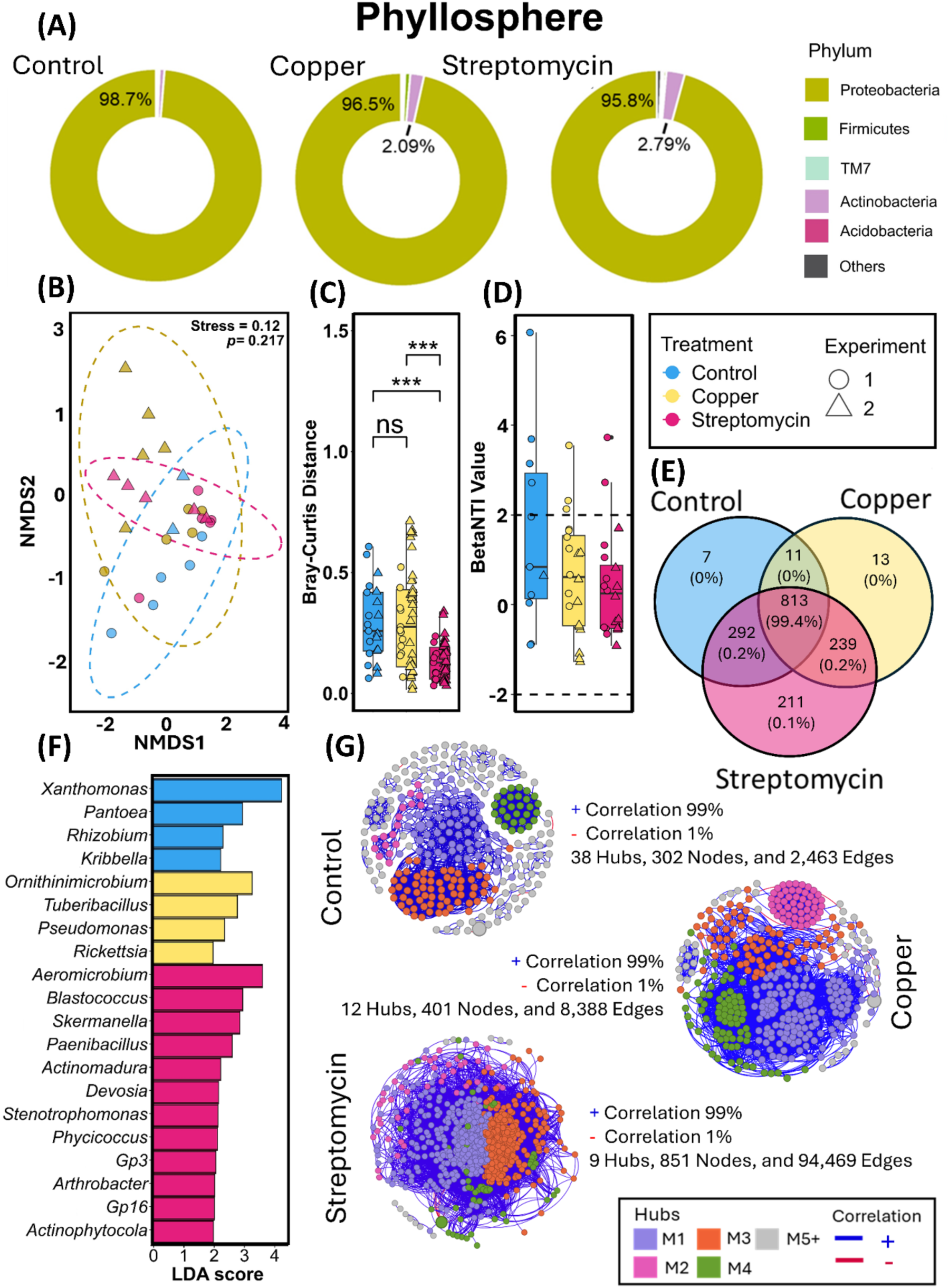
Phyllosphere microbial communities were analyzed for relative taxonomic composition at the phylum level using donut plots (A). Community composition similarity among streptomycin, copper, and control treatments was assessed using NMDS ordination based on Bray-Curtis dissimilarities (B), while Bray-Curtis distance analysis was used to compare treatment separation (C). *β*-nearest taxon index (*β*-NTI) was used to evaluate microbial assembly processes and dysbiosis (D). Venn diagrams identified shared core microbiota, loss of core taxa, and treatment-specific taxa (E). Linear discriminant analysis (LDA) identified differentially enriched taxa among treatments (F), and Spearman co-occurrence networks were used to assess microbial interaction patterns (G).

### Rhizosphere communities remain resilient but are restructured by copper exposure

To assess the robustness of the soil-root interface under antimicrobial influence, we analyzed rhizosphere community dynamics across treatments. Unlike the stochastic volatility shown in aboveground compartments, the rhizosphere communities were comparatively stable across treatments at the phylum level, dominated by Actinobacteria, Proteobacteria, and Acidobacteria (Fig. 4A). NMDS and Bray-Curtis analyses revealed that copper-treated samples exhibited the greatest dispersion, while streptomycin-treated samples clustered tightly and closely resembled control communities. *β*-NTI values were consistently below −2 across treatments, indicating dominance of deterministic assembly processes and limited dysbiosis relative to aboveground compartments (Fig. 4C). A large core microbiome comprising over 8,300 shared OTUs accounted for nearly all community abundance. Copper treatment uniquely increased the number of treatment-specific OTUs and altered network topology, producing expanded but fragmented networks with fewer hubs and a higher proportion of negative correlations, consistent with increased competitive interactions under metal stress.

**Fig. 4.**
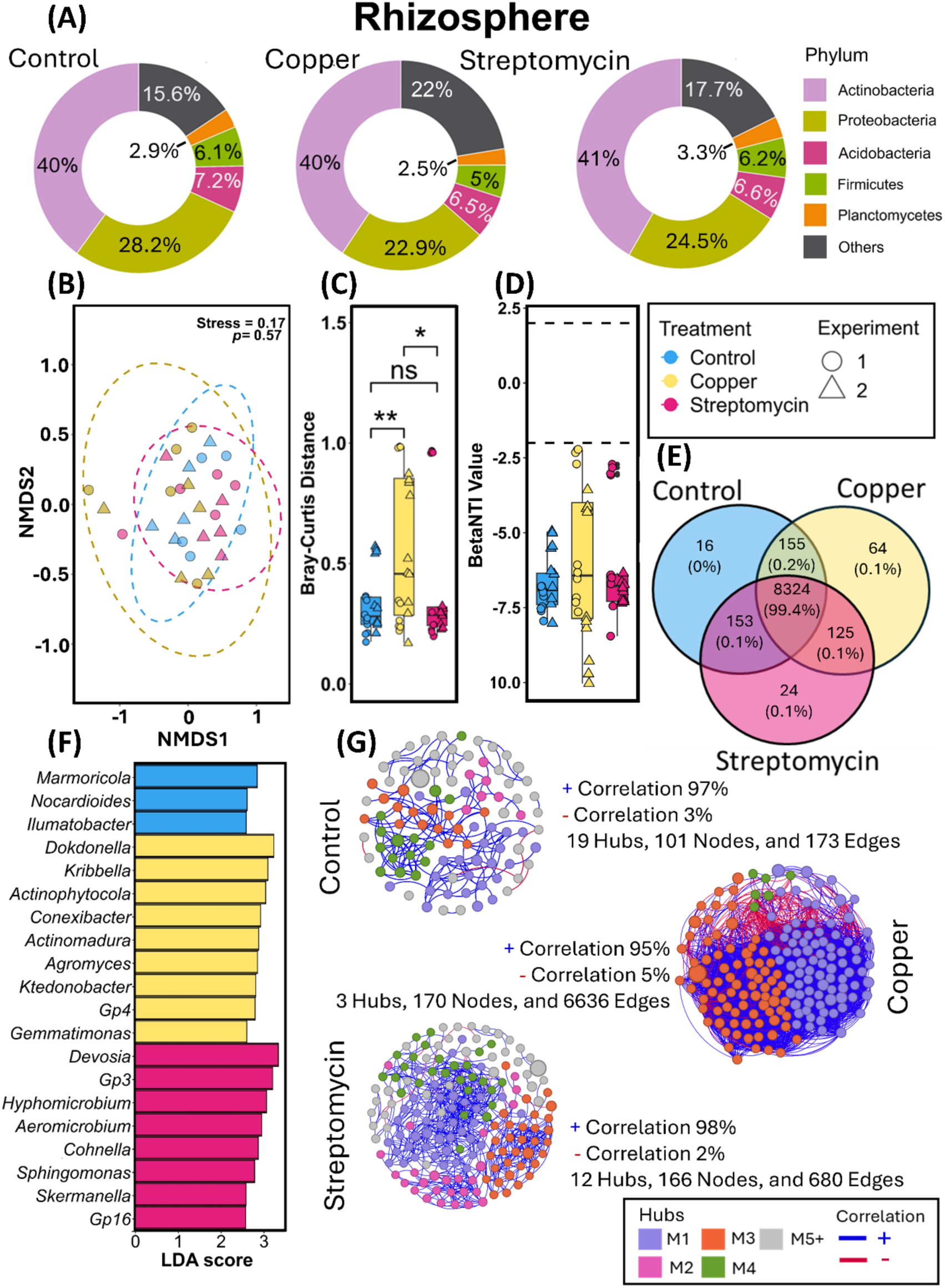
Rhizosphere microbial communities were analyzed for relative taxonomic composition at the phylum level using donut plots (A). Community composition similarity among streptomycin, copper, and control treatments was assessed using NMDS ordination based on Bray-Curtis dissimilarities (B), while Bray-Curtis distance analysis was used to compare treatment separation (C). β-nearest taxon index (*β*-NTI) was used to evaluate microbial assembly processes and dysbiosis (D). Venn diagrams identified shared core microbiota, loss of core taxa, and treatment-specific taxa (E). Linear discriminant analysis (LDA) identified differentially enriched taxa among treatments (F), and Spearman co-occurrence networks were used to assess microbial interaction patterns (G).

### Metagenomics reveals treatment-driven functional restructuring of the microbiome

To examine the rhizosphere functional changes induced by chemical perturbations, gene abundances were compared across control, copper, and streptomycin-treated rhizosphere microbiomes through shotgun metagenomic analyses. Genes showing significant differential abundance (p < 0.05, Log₂ fold change ≥ 1) were assessed. Chemical treatments induced distinct functional reorganizations, reflecting specific selective pressures and differential microbial resilience (Fig. 5). Overall, copper and streptomycin exerted distinct selective pressures on microbiome function. Copper favored enrichment of metal and oxidative stress pathways, whereas streptomycin promoted antibiotic resistance and metabolic adaptation. The concurrent enrichment of microbe-associated molecular pattern (MAMP) associated functions across treatments suggests that chemical dysbiosis may modulate microbial signaling and host interactions. A full list of differentially enriched genes, fold changes, p-values, and predicted taxa is provided in Supplementary Table 1. These findings demonstrate that chemical perturbations drive targeted functional restructuring of plant-associated microbiomes, reflecting both stress-specific selection and broader mechanisms for resilience under dysbiosis.

**Fig. 5.**
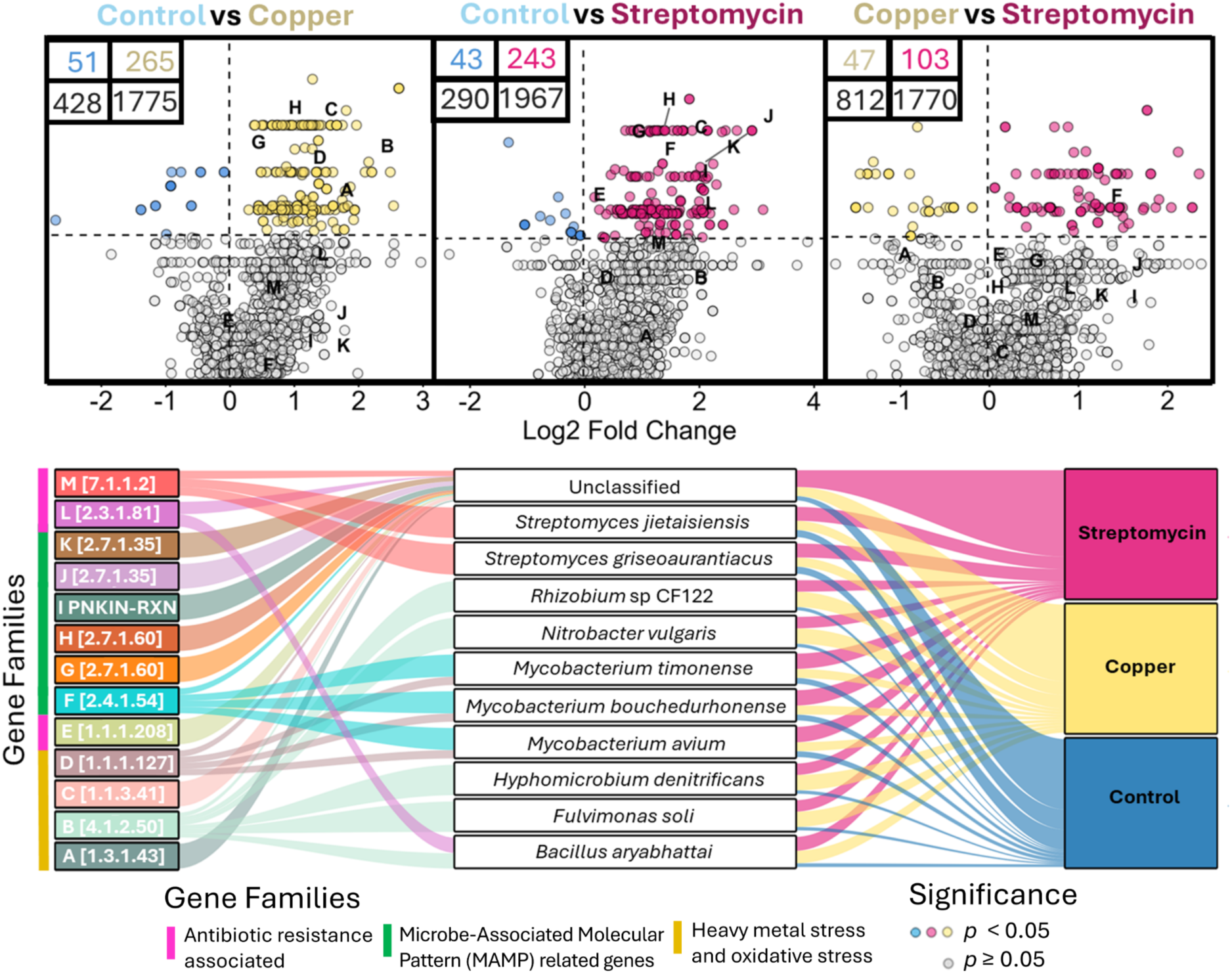
Differential abundance of functional gene families across antimicrobial treatments and their taxonomic origins. (Top) Volcano plots showing differentially abundant gene families for pairwise comparisons: Control vs. Copper (left), Control vs. Streptomycin (center), and Copper vs. Streptomycin (right). Each point represents a gene family, with the x-axis indicating Log₂ fold change and the y-axis showing statistical significance (−log₁₀ adjusted p-value). Dashed lines mark significance thresholds (p < 0.05) and effect size (|Log₂ FC| > 1). Points colored by enriched treatment (blue, yellow, or red) are statistically significant, while gray points are non-significant. Point shading reflects relative abundance. Insets display the number of significant (colored) and non-significant (grey) gene families per quadrant. Letters (A–M) label gene families highlighted in the Sankey diagram below. (Bottom) Sankey diagram showing the relationships between selected gene families (left), their microbial taxonomic origins (center), and the treatment group in which they were enriched (right). Gene families are labeled with EC numbers or pathway identifiers, and colors indicate functional categories: antibiotic resistance related, or heavy metal/oxidative stress response. Ribbon width reflects the relative contribution of each taxon to the abundance of the corresponding gene family within each treatment.

Copper exposure selectively enriched genes linked to metal tolerance and oxidative stress. Notable examples include *6-carboxytetrahydropterin synthase* (Log₂FC=2.14, p=0.010) and *alditol oxidase* (Log₂FC=1.29, p=0.0046), primarily associated with *Bacillus*, *Rhizobium*, and *Mycobacterium*. These functions are consistent with enhanced redox homeostasis and cellular protection under metal stress. Most enrichments were specific to copper-treated communities, suggesting targeted adaptation rather than a generalized stress response. Aditol oxidase was an exception, showing modest enrichment under both copper (Log₂FC=1.29, p=0.005) and streptomycin (Log₂FC=1.63, p=0.005), indicating a broader role in stress resilience.

In streptomycin-treated communities, enrichment of genes linked to antibiotic resistance and energy metabolism was pronounced. *Aminoglycoside N(3)-acetyltransferase*, conferring resistance to aminoglycosides, was significantly enriched (Log₂FC=1.91, p=0.028), primarily encoded by *Bacillus aryabhattai*. Additional genes included *NADH:ubiquinone reductase* (Log₂FC=0.98, p=0.038; associated with *Streptomyces* spp.) and *pyridoxal kinase* (Log₂FC = 2.92, p=0.005), which ranked first among streptomycin-specific enriched genes. These enrichments illustrate selection for antibiotic-resistant and metabolically flexible taxa under dysbiosis.

Across both copper and streptomycin treatments, genes involved in cell wall biosynthesis, amino sugar metabolism, and vitamin B6-dependent redox homeostasis were enriched, including *undecaprenyl-phosphate mannosyltransferase* (Log₂FC=1.55, p=0.005, streptomycin), *N-acylmannosamine kinase* (Log₂FC=1.39, p=0.005, streptomycin; Log₂FC=0.75, p=0.005, copper), and *pyridoxal kinase* (Log₂FC=2.92, p=0.005, streptomycin). These genes support cellular integrity and production of microbe-associated molecular patterns (MAMPs), potentially affecting host-microbe signaling. *Pyridoxal kinase* showed the strongest, treatment-specific enrichment under streptomycin, consistent with a role in oxidative stress mitigation.

## Discussion

The plant microbiome is an integral part of host innate immunity, and disruptions in it increase disease risk^22, 23^. Our study showed that copper- and streptomycin-induced dysbiosis increases disease severity by up to 1.5-fold, suggesting that phyllosphere dysbiosis, similar to that in the rhizosphere, can promote pathogen infection, acting as a disease driver. This observation is consistent with a previous study indicating that disruptions in phyllosphere bacterial communities can influence host susceptibility to pathogens^24^. These results reflect the Anna Karenina Principle, in which stressed microbiomes are more variable and unstable than healthy ones^5^. Our results showed that dysbiosis-inducing antimicrobial treatments, especially streptomycin, together with pathogen infection significantly reduce fruit number, fruit mass, and seed mass, indicating major disruptions in reproductive resource allocation. The effects of copper indicate a gradual decline in microbiome functions, not a total collapse. These findings support evidence that intact native microbiomes are vital for maintaining plant productivity and resilience^25, 26^. The findings from this study suggest that dysbiosis-driven penalties are context-dependent, emerging when human disturbances interact with certain environmental or physiological thresholds. Experiment 1 showed microbiome disruption limits host carbon gain and water use. Streptomycin-induced dysbiosis decreased net CO₂ assimilation (∼1.5 fold) and transpiration (∼1.7 fold) compared to control and copper treatments. These patterns align with frameworks in which immune signaling and stress physiology influence stomatal regulation, potentially reducing carbon gain during defense^27^. Emerging evidence indicates that the microbiome can regulate host physiology through multiple metabolic pathways^28, 29^. For example, previous studies have shown that streptomycin-induced dysbiosis can impair plant physiological function, including reductions in CO₂ assimilation observed after 21 days of antibiotic exposure^6^. Unlike their report of increased intercellular CO₂ (Ci), our results showed no significant change in Ci, implying physiological declines may stem from non-stomatal factors.

A key finding of this study is that identical chemical perturbations elicit compartment-dependent microbiome trajectories rather than uniform responses, with greater instability in transmission-constrained systems such as seeds. Constrained habitats are less redundant and more disturbance-sensitive, leading to varied stability outcomes across plant parts^30^. This niche-specific sensitivity, the maintenance of a eubiotic state in the phyllosphere, is fundamental to gating proper plant immunocompetence, pathogen defense, and stress responsiveness^31, 32, 33, 34^. By contrast, the tighter clustering under copper treatment reflects more sustained deterministic filtering, suggesting a less disrupted community, though likely not a baseline eubiotic state^35^. This interpretation aligns with evidence that microbiome-based plant protection relies more on specific strain identities and interaction patterns within structured assemblages than on community richness^35, 36^.

The seed microbiomes showed the lowest OTU richness across compartments but retained a conserved phylum-level structure dominated by Proteobacteria (syn. Pseudomonadota), consistent with previous findings in tomato and other plant systems^37, 38, 39^. Notably, the relative abundance of Proteobacteria declined from 45% in the control to 43% under copper and 40% under streptomycin, indicating that although this phylum remained dominant, its contribution was reduced under chemical stress. *Xanthomonas* association with streptomycin-treated seeds and *Pseudomonas* enrichment in controls indicate that the treatment mainly caused shifts in dominant taxa, not just in richness^1^. Relative to the control, stress treatments caused major changes in network topology, indicating a restructuring of microbial interactions, consistent with evidence from other studies that simpler microbial networks are less stable and have lower ecosystem functions under stress^40, 41^. This pattern indicates chemical stress reorganizes biotic interactions, reducing ecological complexity^42^. Loss of central hub taxa and connectivity indicates chemical stress filters and weakens the network, limiting core taxa’s role in resilience^22, 43^. The core microbiome, limited to three OTUs, shows a bottleneck in taxon persistence under chemical stress in the seed compartment.

The rhizosphere microbiome exhibited resilience despite copper-induced restructuring, highlighting the importance of root-associated communities in nutrient uptake and stress tolerance ^25, 44, 45^. Unlike the streptomycin-treated phyllosphere, where antibiotics caused severe dysbiosis, the rhizosphere communities exhibited deterministic assembly (*β*-NTI <- 2) and a strong shared core. This shows that belowground microbiomes are mainly influenced by environmental and host factors, even under chemical stress^22, 45^. Moreover, the high diversity of the rhizosphere microbiome facilitates the replacement of lost bacterial taxa through functional redundancy^46, 47^. Within this deterministic framework, copper caused notable changes: increased dispersion, treatment-specific OTUs, and a fragmented network with fewer hubs and more negative correlations. This suggests that metal exposure restructured microbial interactions, promoting competitive filtering and altering ecological interaction^40, 41, 42^. In contrast, streptomycin-treated profiles resembled controls, confirming that antibiotic-driven community destabilization was niche-specific, likely due to the rhizosphere’s superior buffering capacity and distinct assembly mechanisms^44, 48^. From a whole-plant perspective, maintaining an ordered, deterministic rhizosphere assembly, even if restructured, supports host performance and higher fruit yield^22, 25^, whereas the phyllosphere dysbiosis also correlates with the observed disease, physiology and yield penalties^9^.

Copper-driven metagenomic shifts indicate that the treatment induced targeted metal and oxidative-stress responses, favoring microbes that maintain redox balance and metal tolerance over broad stress responses. The enrichment of cofactor- and redox-linked enzymes, such as *6-carboxytetrahydropterin synthase* and *arogenate dehydrogenase*, aligns with evidence that copper-based materials and bactericides alter soil and rhizosphere microbiomes under relevant exposure scenarios^49, 50^. Copper-related changes, including fragmented networks with fewer hubs and more negative correlations, show how metal contamination affects microbial interactions and reorganizes ecological modules^51^. This transition indicates heightened competition among taxa or the collapse of cooperative guilds due to heavy metal pressure, with copper as a key driver of adaptive restructuring rather than community dysbiosis.

Streptomycin-driven enrichment reveals microbiome restructuring, with targeted selection for antibiotic resistance and stress-maintenance determinants^38, 52, 53^. The enrichment of *aminoglycoside N(3)-acetyltransferase* and pathways associated with redox homeostasis (e.g., *NADH:ubiquinone reductase*) and vitamin B6 metabolism (*pyridoxal kinase*) suggest a strong selective pressure under antibiotic exposure. Our results align with previous studies showing that streptomycin-induced dysbiosis increased tomato bacterial spot severity, with host transcriptome reprogramming, reduced metabolism, and increased defense linked to ISR, providing a framework for understanding stressed microbiome states under strong perturbation^6^. These functional shifts align with phenotypic outcomes such as increased disease severity, reduced fruit yield, and decreased gas exchange, consistent with^25, 38, 54^.

Plant pattern recognition receptor (PRR)-mediated recognition is fundamental to pattern-triggered immunity (PTI) and disease resistance^55, 56, 57^. Our shotgun metagenomic results showed environmental stress drives a functional restructuring, focusing on cell-envelope remodeling, amino-sugar metabolism, and redox homeostasis, which can occur even when overall community colonization levels remain relatively constant^34, 58, 59^. This dysbiosis-driven transition toward microbial stress-survival and envelope-remodeling functions may heighten physiological costs and weaken host immunity, underscoring the need for a healthy microbiome for plant defense^6, 31^. The streptomycin-specific enrichment of pyridoxal kinase and undecaprenyl-phosphate mannosyltransferase suggests that microbes prioritize redox balance and envelope biogenesis to combat antibiotic stres^60, 61^.

Across compartments, microbiome dysbiosis altered taxonomic composition, increased community dispersion, modified ecological assembly processes, and reshaped network connectivity, with the magnitude and nature of these effects varying among seed, phyllosphere, and rhizosphere environments. These findings emphasize the importance of microbiome stability in maintaining plant health under stress.

## Methods

### Plant material and soil source

Experiments were conducted using the tomato plant (*Solanum lycopersicum* L.) cultivar Alisia Craig as a model plant for dysbiosis studies. Soil was collected from a field with a history of organic solanaceous crop cultivation (potato, tomato, and sweet potato) at the University of Florida, USA (elevation 26 m; 29°38’14.0832” N, 82°21’40.6238” W). Prior to sowing, physicochemical properties were characterized to enable robust interpretation of microbiome dynamics and disease outcomes (Table S2). Field soil was mixed with a commercial potting substrate containing Canadian sphagnum peat and perlite (Jolly Gardener HFC 20), in a 1:1 ratio, following ^6^, to maintain suitable soil moisture in the experiment

Seeds were sown in 2.0 L pots, and plants were maintained in a controlled greenhouse environment at the University of Florida (29°38’23.7305” N, 82°21’28.3402” W) under standard conditions adjusted for 27-32 °C and 70-85% relative humidity. At 3 weeks old, five seedlings were allocated to each treatment. Plants were watered daily after inoculation until field capacity. Six seedlings were assigned to each treatment group.

### Rhizosphere and phyllosphere dysbiosis induction and pathogen inoculation

To induce dysbiosis, tomato plants were divided into three treatment groups. The first group received only sterile distilled water (T1 - Control), the second group received copper hydroxide (Kocide 3000 Mitsui Ltd Houston, TX, USA) at 1.5 g L^-1^ (T2 - Copper), and the third group received streptomycin sulfate (Fisher Scientific Co LLC, USA) at 0.6 g L^-1^ (T3 - Streptomycin). For all groups, the respective solutions were prepared in distilled water and applied at two different developmental stages. A total of 54 tomato plants were used in the experiment (three treatments x six repetitions x three plant parts – rhizosphere, phyllosphere, seeds), and the experiment was repeated twice.

Tomato plants were first inoculated with *Xanthomonas perforans* at 21 days after emergence using a spray inoculation method (1 × 10⁵ cells mL⁻¹) until runoff. The inoculum was prepared from a single pure colony of *Xanthomonas perforans* (streptomycin-resistant strain XP1-6) grown in lysogeny broth (LB) at 28°C with agitation (150 rpm) for 24 h. Cells were collected by centrifugation (10,000 × g, 10 min, 25°C), resuspended in sterile distilled water, and adjusted to an OD600 of 0.1. Immediately after inoculation, plants were covered with translucent polypropylene bags for 24 h to maintain humidity and promote infection establishment.

Following pathogen inoculation, treatments were applied to evaluate the microbiome state of dysbiosis across plant compartments. For rhizosphere dysbiosis, treatments were applied once, whereas phyllosphere treatments were applied twice. The first application consisted of a rhizosphere soil drench with 200 mL of treatment solution per plant. A second application was performed at flowering as a spray, when approximately 90% of plants had developed 1–2 open flowers. At this stage, treatments were applied to the phyllosphere by spraying the canopy to runoff using a manual 1.5 L sprayer. This sequential treatment design was intended to reinforce dysbiosis induction in both below-ground and above-ground microbiomes following pathogen establishment, as shown in (Fig. 6).

**Fig. 6.**
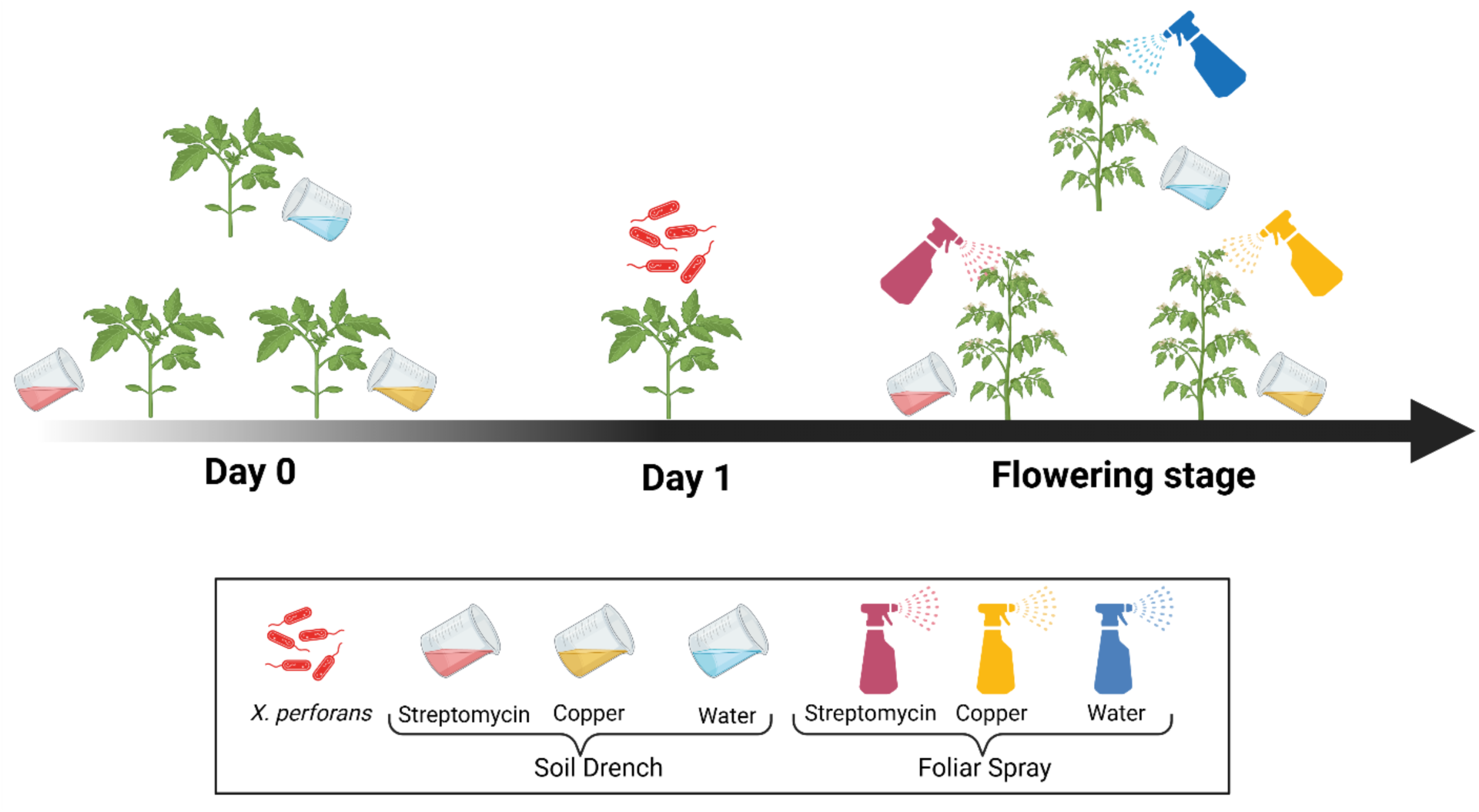
Experimental timeline of treatment application and pathogen challenge in tomato. Streptomycin, copper hydroxide or water were applied to the rhizosphere at 21 days after emergence (day 0) and 24 h later *Xanthomonas perforans* was inoculated (day 1). At flowering stage, treatments were reapplied to the rhizosphere and phyllosphere via soil drenching and foliar spraying, respectively.

### Disease severity, yield, and physiological measurements

Disease assessments began at 5 days after inoculation (5 DAI) and were repeated every 3 days, for a total of seven evaluations. Bacterial spot severity on each plant was quantified using the Standard Area Diagram (SAD) for tomato leaf bacterial spot, which provides 12 reference images ranging from 0.5% to 90% of leaf area^62^. The Bacterial Spot (BS) was assessed on a set of six leaflets identified in each true leaf, with petioles marked for each plant. The percentage severity of diseased leaf area was visually estimated by matching symptoms to the closest severity class on the SAD scale. Both necrotic and chlorotic symptoms were included in the total disease severity score, in accordance with the criteria established in the SAD. Disease progress over time was summarized by calculating the Area Under the Disease Progress Curve (AUDPC) as described in^63^.

To evaluate the effects of copper hydroxide and streptomycin-induced dysbiosis on agronomic performance, the plant growth and yield-related traits were assessed. Plant fresh weight was measured by cutting each plant about 2 cm above the soil surface and weighing the entire aerial Each plant was then placed in a paper bag after its fresh weight was recorded on an analytical balance (ATX224, Shimadzu Corporation; Kyoto, Japan). Next, the plant shoots were transferred to a forced-air drying oven and dried at 70 °C until reaching a constant weight. The dry mass of the plant’s aerial parts was recorded to calculate the overall dry biomass. Fruits were harvested continuously as they ripened until 100 days post-transplanting. Total fruit yield was assessed as the number of fruits per plant and the mean fresh mass per fruit, measured using an analytical balance (ATX224, Shimadzu Corporation; Kyoto, Japan). For seed mass determination, seeds were aseptically extracted under a laminar flow hood and air-dried on sterile filter paper for 48 h at room temperature. Seed mass was determined by weighing 25 seeds per treatment and replicate on an analytical balance, thus calculating the mean seed mass.

The physiological parameters were assessed at 24 DAI, using measures of gas exchange and photosynthesis rate. Observations were conducted with an infrared gas analyzer (IRGA) LI-COR portable photosynthesis system (LI-6800, Li-Cor, Lincoln, NE, USA). Five plants/treatments were randomly selected to assess photosynthesis, gas exchange, stomatal function, and transpiration characteristics. Parameters: flow, water, and CO2 injector: activated; humidifier: 20%; CO2 concentration: 20%; CO_2_ reference: 420 mmol; H_2_O reference: 65%; desiccant: 100%; relative humidity: 50%; light intensity: 1000 mmol, 9:1 red/blue ratio; temperature: 27°C. Three measurements (technical replicates) were conducted on hardened leaves per plant, which were characterized by a darker green hue, a glossy texture, and firmness, suggesting full development and optimal photosynthetic capacity.

### Tomato rhizosphere, phyllosphere, and seed microbiome extraction

For the rhizosphere and phyllosphere, samples were collected 24 days after pathogen inoculation (DAI). For rhizosphere microbiome extraction, approximately 7 g of the distal root system, including tightly adhering soil, was collected from the distal root portions. Roots were excised and transferred to a sterile 50 mL polypropylene tube containing 25 mL of potassium phosphate buffer (6.75 g of KH_2_PO_4_, 8.75 g of K_2_HPO_4_, and 1 mL of Triton X-100 in 1 L of distilled water), called removal buffer A, then immediately placed on ice. The tubes containing rhizosphere soil were vortexed for 30 seconds and subsequently sonicated in an ultrasonic bath for 10 min at room temperature, following a previously described protocol with minor modifications^64^. After sonication, roots were aseptically removed from the tubes and discarded. The remaining suspension was centrifuged at 4 °C and 4,000 × g for 10 minutes, and the supernatant was carefully discarded. The resulting pellet, representing the rhizosphere microbiome, was stored at -80 °C until DNA extraction.

Phyllosphere microbiome extraction followed the protocol outlined by^65^, with some adjustments. For each sample, 18 leaflets were randomly selected, transferred to a sterile 50 mL centrifuge tube, and submerged in 25 mL of potassium phosphate buffer. This buffer had the same composition as removal buffer A, except that Triton X-100 was replaced with 0.05% Tween 20 (Sigma-Aldrich, Burlington, MA, U.S.A.), called removal buffer B. Tubes were placed in an ultrasonic bath (VEVOR-Ultrasonic Cleaner, Guangdong, China) and sonicated for 15 minutes at room temperature, then vortexed for 20 seconds. After the first sonication, leaflets were aseptically transferred to a second sterile centrifuge tube with 25 mL of potassium phosphate buffer plus 0.05% Tween 20, and the sonication and vortex steps were repeated once as suggested by^66^. After the second extraction, all leaf tissue was discarded. Both suspension tubes were centrifuged at 7,197 × g for 2 minutes to pellet the microbial cells. The supernatant from each tube was carefully removed, leaving about 1 mL of liquid with the pelleted cells. The cell pellets were resuspended by vortexing and pooled into a single tube, resulting in approximately 2 ml. The cell pellets were stored at -80 °C until further DNA extraction.

For seed microbiome extraction, only fully mature, ripe tomato fruits were used as seed sources. Seed extraction was performed aseptically with sterile scalpels and forceps inside a laminar flow cabinet to prevent external contamination. The seeds were then air-dried on sterile filter paper at room temperature under aseptic conditions. Afterward, 24 individual seed samples were sliced in half with a sterile razor blade to enhance endophytic microbial release, as described by^67^. Thus, the seeds were processed according to the methods described by ^1^, with minor adjustments. Seeds were immersed in removal buffer B at a ratio of 2 mL buffer per gram of seed. They were soaked at 20 °C on a shaker operating at 150 rpm for 2 hours and 30 minutes. Seed tissues were carefully removed with sterile forceps, and the resulting suspension was centrifuged at 10,000 × g for 10 minutes to pellet microbial cells. The supernatant was gently discarded, and the pellet containing the seed-associated microbiome was stored at -80 °C until DNA extraction.

### DNA extraction, quality control, sequencing and microbiome bioinformatic analysis

Genomic DNA was extracted from 200 mg per sample of the rhizosphere fraction measured as net mass and the phyllosphere fraction (wet mass), and from 100 mg per sample of the seed microbiome fraction (wet mass). The DNA was extracted using the Zymo Quick DNA Faecal/Soil Microbe MiniPrep Kit (Zymo Research, Irvine, CA, USA) according to the manufacturer’s instructions. DNA concentration and quality were assessed using a NanoDrop spectrophotometry (Thermo Fisher). The DNA was dispatched for sequencing (rhizosphere, *n* = 10; phyllosphere, *n* = 10; seed, *n* = 10) to SeqCenter (Pittsburgh, PA; seqcenter.com). The 16S rRNA gene in the V3-V4 regions was sequenced. Following purification and normalization, the samples were sequenced using a V3 MiSeq 622 cycle flow cell, producing 2 × 301 bp paired-end reads.

For rhizosphere samples (n = 10), an additional aliquot of genomic DNA was submitted for whole-metagenome shotgun sequencing at SeqCenter (Pittsburgh, PA, USA). Sequencing was performed on an Illumina NovaSeq X Plus platform using short-read paired-end chemistry, generating 2 × 151 bp reads. Libraries were prepared with the Illumina DNA Prep kit using IDT 10 bp unique dual indices and a target insert size of 280 bp, with no additional fragmentation or size selection. Demultiplexing, quality control, and adapter trimming were carried out using bcl-convert v4.2.4.

### Statistical analysis of phenotypic and physiological traits

Agronomic, disease severity and physiological data analyses were performed in R version 4.5.1. Each experimental unit consisted of three plants. Within each block, one plant per plot was used to measure plant fresh and dry mass and to sample the rhizosphere and phyllosphere microbiomes, whereas the remaining two plants were used for yield measurements, including number of fruits per plant, fruit fresh mass, number of seeds per fruit, and seed mass. Disease severity ratings were converted to the area under the disease progress curve (AUDPC). Response variables included AUDPC, plant fresh mass, plant dry mass, number of fruits per plant, fruit mass, number of seeds per fruit, seed mass, and gas exchange variables measured by infrared gas analysis.

Data were analyzed using linear mixed-effects models. For each response variable, an initial model was fitted including treatment, experiment, and their interaction as fixed effects, and block nested within experiment as a random effect. When the treatment × experiment interaction was not significant (p > 0.05), data were combined; otherwise, experiments were analyzed separately. Heteroskedastic mixed-effects models were then fitted to address potential heterogeneity in residual variances across treatments, using the nlme package and specifying treatment-specific residual variances^68^. Final models included Treatment and Experiment as fixed effects, Block within Experiment as a random effect, and residual variance structures that depend on treatment. Estimated marginal means (EMMs) and treatment-specific standard errors (SEs) were calculated using the emmeans package^69^. Pairwise comparisons among treatments were performed using Tukey’s honestly significant difference (HSD) test, applying correction for multiple comparisons. All graphics were created in R with ggplot2, showing treatment means with SE bars from the fitted models, raw block-level data as jittered points.

### Statistical analysis of microbial communities based on 16S rRNA gene sequencing

Microbial community analyses were conducted using amplicon sequence data generated from OTU tables in Mothur using Miseq SOP^70^. Alpha diversity metrics, including Shannon diversity, observed richness, and Pielou’s evenness, were calculated using the estimate_richness function from the phyloseq package^71^. To assess differences in alpha diversity among treatment groups, pairwise comparisons were performed using Wilcoxon rank-sum tests. *P*-values were corrected for multiple testing using the Benjamini-Hochberg false discovery rate (FDR) procedure.

Beta diversity was quantified using Bray-Curtis dissimilarity matrices, and differences in community composition among treatments were visualized through non-metric multidimensional scaling (NMDS) implemented in the vegan R package^72^. Permutational multivariate analysis of variance (PERMANOVA) was applied using the adonis2 function to test for statistically significant differences in overall community structure among treatments. To further identify pairwise differences between treatments, Bray-Curtis distances were compared using Wilcoxon rank-sum tests with Benjamini-Hochberg FDR correction. Phylogenetic turnover among microbial communities was quantified using the beta-nearest taxon index (betaNTI) calculated from CSS-normalized OTU tables. *β*-NTI was computed with 999 null model iterations and abundance weighting using the trans_nullmodel package, employing both “sample.pool” and “taxa.labels” null models. Resulting *β*-NTI values were used to calculate group-level distances with trans_beta, and visualized as boxplots for each treatment, with individual experiments distinguished by point shape. Dashed lines at |betaNTI| = 2 indicate thresholds for deterministic versus stochastic community assembly.

Composition similarities among treatment groups (control, copper hydroxide, and streptomycin) were evaluated using Venn diagrams to visualize shared and unique OTUs. To identify taxa associated with specific treatments, linear discriminant analysis (LDA) was performed with a relaxed significance threshold (α = 0.5) to detect more potential discriminative organisms. Microbial co-occurrence networks were constructed separately for each treatment using filtered OTU tables. Networks were inferred based on Spearman correlations, with significant correlations defined as p < 0.05, and optimized for sparsity. Network structures and modules were visualized and explored using Gephi v0.10.1^73^.

### Shotgun metagenomic analysis

Raw sequences were evaluated for quality using FastQC. Quality and adapter trimming were then performed within the University of Florida hypergator High performance computer using trim_galore v. 0.2.8 using default settings and cut adapt v.1.18 ^74^. The success of the quality filtering was confirmed by a subsequent FastQC analysis. After passing quality control merged FASTA files the functional profiling was performed using HUMAnN v3.8 and Uniref database to identify “genefamilies” from our datasets. The composition of microbial communities for eight datasets was computed using the MetaPhlAn version 4.0.2.^75^. Resulting gene families were annotated with their corresponding organism from the HUMAnN and MetaPhlAn results.

Gene abundance data were analyzed to identify treatment-associated differences. Only genes detected in at least two treatments with ≥2 replicates per treatment were included, using tidyr and dplyr in R^76, 77^. Differential abundance was assessed with the non-parametric Mann-Whitney U test and the parametric two-sample t-test, and effect sizes were estimated as Cohen’s d (stats package)^69^. Results were compiled into structured tables summarizing all genes, comparison-specific outcomes, and significant genes. Differentially abundant genes were visualized using volcano plots generated with ggplot2^78^, with log₂ fold changes on the x-axis and −log₁₀-transformed Mann-Whitney p-values on the y-axis. Gene families of interest were identified based on statistical significance (Mann-Whitney p < 0.05) and known associations with microbe-associated molecular patterns (MAMPs), copper/heavy metal stress, or antibiotic resistance. Associations between microbial gene families, bacterial taxa, and treatments were visualized using a custom Sankey diagram implemented in Python (pandas, matplotlib), with ribbon widths proportional to gene abundance to illustrate the directional flow from defense-related enzymes to microbial species and treatment conditions^79, 80^.

## Supporting information

Supplementary Doc 1

## Acknowledgements

This work was supported by the USDA National Institute of Food and Agriculture (NIFA) project no. 2022-68015-36721, the University of Florida’s Institute of Food and Agricultural Sciences (UF*/*IFAS) seed grant, and by the Research Capacity Fund (Hatch) program, project award no. 7010682.

## Author contributions

Conceptualization: SJM, EMG, FJP, and TK; investigation: FJP, TK, and MRT; data analysis and statistical methodology: VET, Figure preparation: VET, data curation: VET, FJP, and SMT; writing-review and revision: VET, FJP, SJM, EMG, TK, SDM; project administration, SJM and EMG. All authors have read and agreed to the published version of the manuscript.

