## Supplementary Doc 1 for "Antibiotics and copper drive compartment-specific dysbiosis and functional reprogramming in tomato microbiomes"

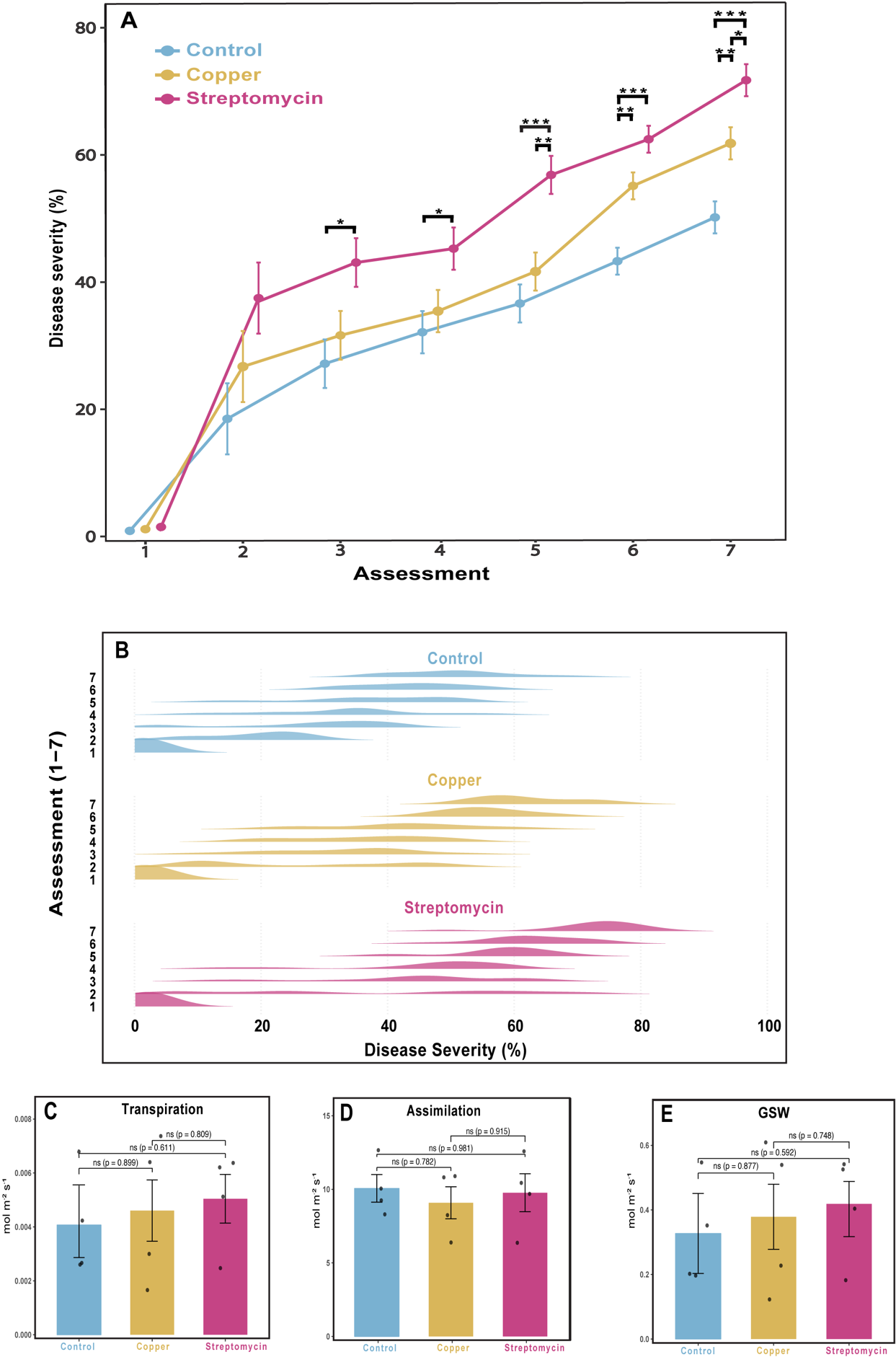

**Figure S1. (A)** Disease progress curves showing bacterial spot severity (%) caused by *Xanthomonas perforans* in tomato plants. Disease severity was monitored over seven assessments in plants treated with Control, Copper, or Streptomycin. Curves represent estimated marginal means (EMMs) pooled across two independent experiments, and error bars indicate SE. Asterisks denote significant pairwise differences between connected treatments based on Tukey’s post hoc multiple comparisons test (*P < 0.05, **P < 0.01, *P < 0.001). Streptomycin showed higher disease severity than Control from assessments 3-7 and Copper from assessments 5–7, whereas Control exceeded Copper at assessments 6-7. **(B)** Ridgeline density plots showing the distribution of bacterial spot severity (%) in tomato plants across seven assessments under Control, Copper, or Streptomycin treatments. Distributions pooled from two independent experiments showed that severity increased over time across all treatments, with Streptomycin highest, followed by Copper, and Control remaining lowest. **(C-E)** Physiological responses of tomato plants in Experiment 2 under Control, Copper and Streptomycin treatments. (C) Transpiration, (D) net CO₂ assimilation and (E) stomatal conductance (gsw). Bars represent means ± SE. No significant differences were detected among treatments for any of the physiological parameters, based on Tukey’s post hoc multiple comparisons test (ns, not significant).

**Supplementary Table S1 | Soil chemical properties relevant to microbiome analyses.** Soil sample analyzed by the UF/IFAS Analytical Research Laboratory.

| ***Parameter*** | pH | Total *N* | Total *P* | *P* | *K* | *Ca* | *Mg* | *S* | *Al* | *Fe* | *Mn* | *Zn* | *Cu* | *B* |
| --- | --- | --- | --- | --- | --- | --- | --- | --- | --- | --- | --- | --- | --- | --- |
| ***Value*** | 6.60 | 0.09 | 756.82 | 337.62 | 205.95 | 1611.97 | 202.36 | 163.72 | 610.00 | 155.69 | 30.99 | 11.23 | 1.74 | 0.60 |
|  | --- | % | *mg kg⁻¹* | | | | | | | | | | | |
